# SARS-CoV-2 T cell responses are expected to remain robust against Omicron

**DOI:** 10.1101/2021.12.12.472315

**Authors:** Syed Faraz Ahmed, Ahmed Abdul Quadeer, Matthew R. McKay

**Affiliations:** Department of Electronic and Computer Engineering, The Hong Kong University of Science and Technology, Clear Water Bay, Hong Kong SAR, China; Department of Chemical and Biological Engineering, The Hong Kong University of Science and Technology, Clear Water Bay, Hong Kong SAR, China; Department of Electrical and Electronic Engineering, University of Melbourne, Melbourne, Victoria, Australia; Department of Microbiology and Immunology, University of Melbourne, at The Peter Doherty Institute for Infection and Immunity, Melbourne, Victoria, Australia

## Abstract

Omicron, the most recent SARS-CoV-2 variant of concern (VOC), harbours multiple mutations in the spike protein that were not observed in previous VOCs. Initial studies suggest Omicron to substantially reduce the neutralizing capability of antibodies induced from vaccines and previous infection. However, its effect on T cell responses remains to be determined. Here, we assess the effect of Omicron mutations on known T cell epitopes and report data suggesting T cell responses to remain broadly robust against this new variant.

## MAIN TEXT

Omicron (B.1.1.529), detected in Botswana and South Africa in early November 2021, has emerged as a new SARS-CoV-2 variant of concern (VOC). Based on SARS-CoV-2 genome data available on GISAID (https://www.gisaid.org), Omicron has been identified in at least 60 countries (as of 10 December 2021), raising concerns about its potentially high transmissibility. The most worrying aspect of Omicron is the abundance of mutations in the spike (S) protein, with some shared with previous VOCs and some new. Preliminary studies have reported a drastic reduction in the neutralization efficacy of infection- and vaccine-elicited antibodies and sera against Omicron (Wilhelm et al. 2021; Roessler et al. 2021), indicating a strong capability of Omicron to evade humoral immune responses. However, the extent to which Omicron is capable to evade cellular immune responses, the other arm of the adaptive immune system mediated by T cells, is not yet clear.

T cell responses are a key armament against viral infections which, in addition to assisting B cell activation for generating antibodies, help in providing protection from disease by eliminating virus-infected cells. SARS-CoV-2 T cell responses induced by either natural infection or vaccines have been linked to rapid viral clearance and reduced disease severity (Rydyznski Moderbacher et al. 2020; Kalimuddin et al. 2021; Bertoletti, Tan, and Le Bert 2021), even when the neutralizing antibody response is reduced (Geers et al. 2021) or absent (Steiner et al. 2021). Thus, if SARS-CoV-2 T cell responses hold up, they are likely to assist in limiting disease severity in infections caused by Omicron that seemingly escapes neutralizing antibodies (Wilhelm et al. 2021; Roessler et al. 2021). However, if mutations in Omicron result in T cell escape, it could limit the protection provided by T cells.

Here we provide a preliminary investigation into the robustness of T cell responses against Omicron by leveraging information of T cell epitopes known to be targeted in COVID-19 infected and/or vaccinated individuals. We first focused on epitopes derived from the S protein. This is of primary interest in the context of T cell responses elicited by COVID-19 vaccines, with many such vaccines employing S-specific antigens. T cell responses against S have also been shown to be immunodominant upon natural infection (Tarke, Sidney, Kidd, et al. 2021). We considered all S-specific 224 CD8^+^ and 167 CD4^+^ SARS-CoV-2 T cell epitopes available at IEDB (Vita et al. 2019) and screened them for Omicron-defining mutations (obtained from https://covariants.org). This revealed that 14% of CD8^+^ and 28% of CD4^+^ T cell epitopes comprise at least one position harbouring an Omicron mutation (**Figure 1**), indicating that a large majority of both CD8^+^ and CD4^+^ T cell epitopes still appear to remain unaffected by Omicron.

**Figure 1.**
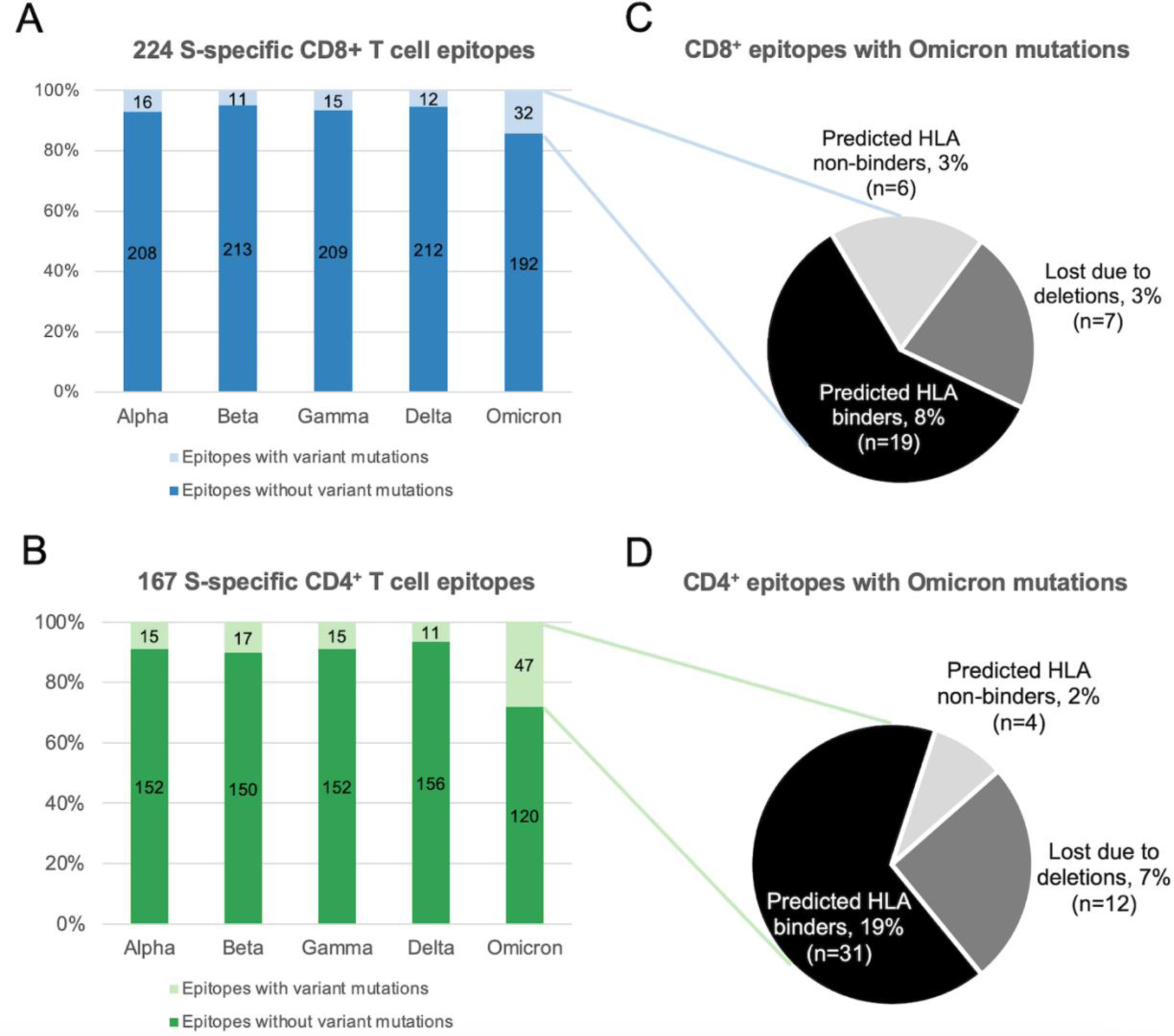
(**A**-**B**) Percentage of S-specific SARS-CoV-2 T cell epitopes with and without mutations present in the five VOCs: (**A**) CD8^+^ T cell epitopes. (**B**) CD4^+^ T cell epitopes. Only epitopes of canonical lengths (CD8^+^: 8-12 residues and CD4^+^:15 residues) were included in the analysis. VOC-defining mutations (that also include deletions) were obtained from https://covariants.org. (**C**-**D**) Predicted effect of Omicron mutations on peptide-HLA binding of SARS-CoV-2 (**C**) CD8^+^ and (**D**) CD4^+^ T cell epitopes. NetMHCpan-4.1 and NetMHCpanII-4.0 were employed for predicting peptide-HLA binding using the default parameters (Reynisson et al. 2020).

The number of epitopes encompassing Omicron mutations (32 CD8^+^ and 47 CD4^+^ epitopes) are nonetheless notably higher than for other VOCs, especially in the case of CD4^+^ T cell epitopes. To further assess the capacity of Omicron mutations to evade responses against these epitopes, we employed widely-used computational tools (Reynisson et al. 2020) to predict the impact of Omicron mutations on binding of these S-specific epitopes to their cognate HLA alleles. Such HLA-epitope binding is a necessary requirement for T cell recognition. Inspection of these mutations revealed that the multiple deletions in the S protein of Omicron lead to loss of seven CD8^+^ and twelve CD4^+^ epitopes, while only six CD8^+^ and four CD4^+^ epitopes (listed in **Supplementary Tables 1 and 2**, respectively) are predicted to become non-binders (**Figures 1C-D**). In the case of CD8^+^ epitopes, half (3/6) of the epitopes predicted to abrogate HLA binding are exclusive to Omicron, while the remaining were also observed in previous VOCs (**Supplementary Table 1**). Since the majority of the CD8^+^ and CD4^+^ epitopes harbouring Omicron mutations are predicted to retain HLA binding (**Figures 1C-D**), this lowers the possibility of T cell escape.

Taken collectively, the presented data would suggest that SARS-CoV-2 T cell immunity acquired by S-focused COVID-19 vaccines or previous infection would remain broadly robust against Omicron, as was the case for other VOCs (Stanojevic et al. 2021; Tarke, Sidney, Methot, et al. 2021).

As an interesting by-product, our analysis revealed that a new peptide EPEDLPQGF, gained for the first time due to the unique three amino acid insertion ‘EPE’ following position 214 in Omicron, is predicted to bind strongly with two common HLA alleles (HLA-B*35:01 and B*53:01) and weakly with four others (HLA-A*26:01, B*07:02, B*44:02, and B*51:01). This peptide may constitute a unique Omicron-specific T cell epitope, if confirmed to be targeted.

We next considered SARS-CoV-2 T cell epitopes derived from other proteins, besides S. This is of interest in the context of T cell responses induced by prior natural infection or inactivated COVID-19 vaccines, which have been found to target epitopes derived from multiple SARS-CoV-2 proteins (Grifoni et al. 2020; Deng et al. 2021). Investigating all 745 CD8^+^ and 373 CD4^+^ SARS-CoV-2 T cell epitopes derived from proteins besides S (available at IEDB (Vita et al. 2019)) revealed that an overwhelming majority of them (98% and 97% respectively) remained unaffected by mutations observed in Omicron (**Figure 2**). Among those affected, deletions accounted for the putative loss of five epitopes, with four (1 CD8^+^ and 3 CD4^+^) being from the immunodominant nucleocapsid (N) protein.

**Figure 2.**
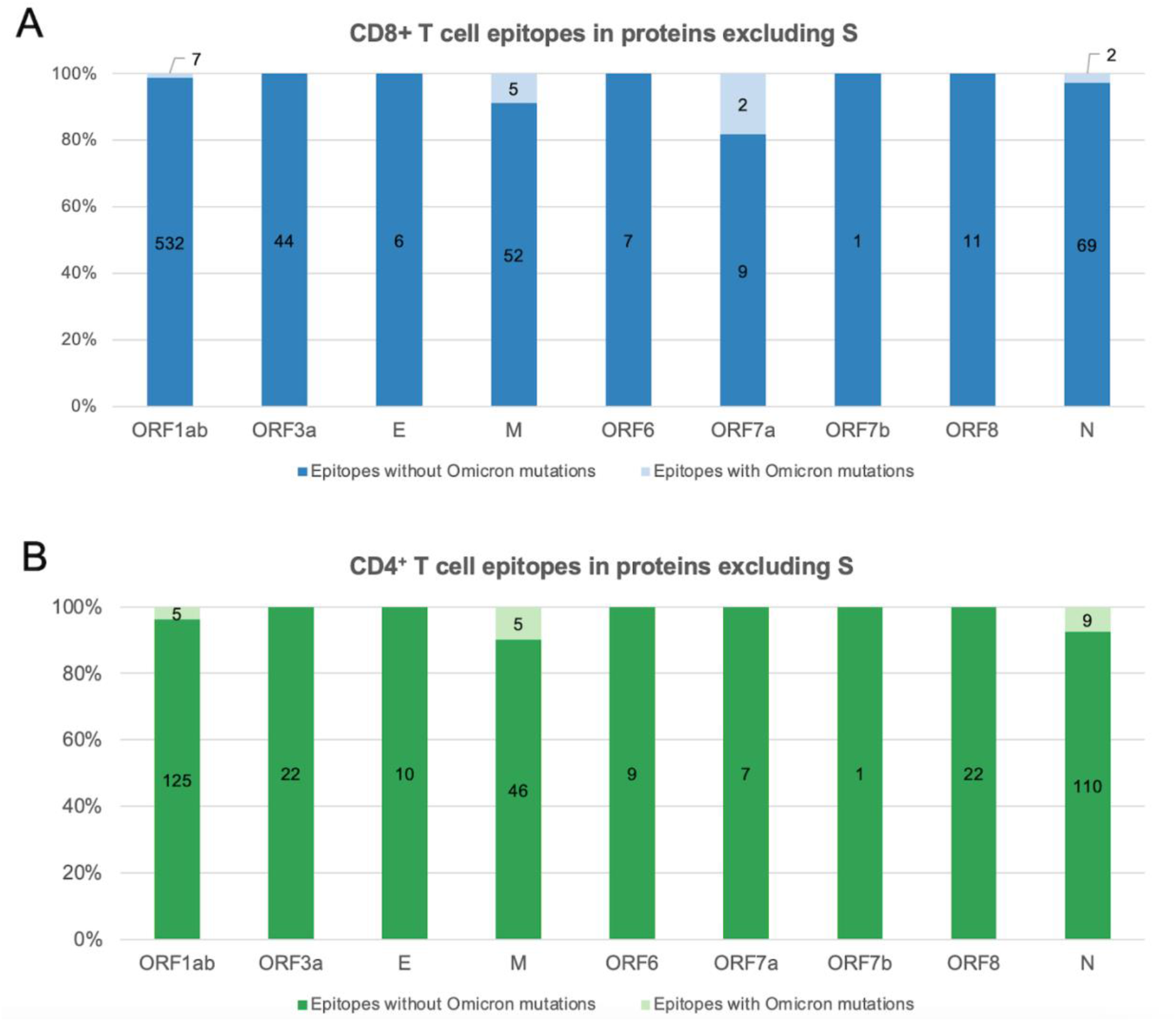
Percentage of SARS-CoV-2 T cell epitopes derived from proteins besides S with and without Omicron-defining mutations: (**A**) CD8^+^ T cell epitopes. (**B**) CD4^+^ T cell epitopes. Only epitopes of canonical lengths (CD8^+^: 8-12 residues and CD4^+^:15 residues) were included in the analysis. Omicron-defining mutations (that also include deletions) were obtained from https://covariants.org.

It is the case that some T cell epitopes are preferentially targeted in the population. Escape from responses against the most commonly targeted epitopes should be carefully scrutinized, since it could potentially affect a large fraction of the population. To investigate this, we assessed whether Omicron mutations affect immunoprevalent CD8^+^ T cell epitopes that have been reported to be commonly targeted by convalescent individuals from different geographical regions (Quadeer, Ahmed, and McKay 2021). As was the case for other VOCs, none of the Omicron mutations were present in any of the 20 immunoprevalent epitopes. Coupling these results with the fact that multiple T cell epitopes are targeted within an individual (Tarke, Sidney, Kidd, et al. 2021), suggests that the acquired T cell immunity across proteins would be expected to remain largely intact against Omicron.

There are multiple limitations of this study. First, the study is based on a set of experimentally-determined SARS-CoV-2 T cell epitopes that have been reported so far. While this set includes a large number of epitopes, it may not be exhaustive. Second, assessing the robustness of T cell responses against a SARS-CoV-2 variant using the fraction of epitopes encompassing mutations present in the variant is a first order analysis. While it can assist to provide preliminary estimates, further targeted experiments are required to confirm the robustness of T cell responses against Omicron, and also to test the capacity of specific epitope mutations to confer T cell escape. Nevertheless, given that most of the experimental T cell epitopes known to be targeted in vaccinated and/or previously infected individuals (collectively, ∼60% of the global population (Hannah Ritchie Edouard Mathieu and Roser 2020)) are unaffected by Omicron mutations, our preliminary analysis suggests that the effectiveness of pre-existing T cell immunity will remain intact. Thus, while Omicron might lead to an increase in the number of infections due to increased transmissibility and antibody escape, robust T cell immunity provides hope that, similar to other VOCs (UK-Health-Security-Agency 2021), the level of protection against severe disease would remain high.

## ACKNOWLEDGMENTS

This analysis was made possible by the open sharing of immunological data of experimentally determined SARS-CoV-2 epitopes by research groups from around the world through the IEDB database. We gratefully acknowledge the contributions of all the researchers, scientists and technical staff involved. We would also like to thank the service of South African research teams involved in the early detection of Omicron, without which the world would not have known about it. This work was supported by the General Research Fund of the Hong Kong Research Grants Council (RGC) (Grant no. 16213121).

## COMPETING INTERESTS

The authors have filed for patent protection for various applications of SARS-CoV-2 T cell epitopes.

## SUPPLEMENTARY TABLES

**Supplementary Table 1.** List of CD8^+^ T cell epitopes with Omicron mutations that are predicted to become non-binders. NetMHCpan-4.1 was employed for predicting peptide-HLA binding using the default parameters (Reynisson et al. 2020).

| No. | Epitope | HLA | Mutation(s) | VOC |
| --- | --- | --- | --- | --- |
| 1 | FCNDPFLGVYY | A*01:01 | Y145D | Omicron |
| 2 | YYHKNNKSW | A*24:02 | Y145D | Omicron |
| 3 | YGFQPTNGV | B*51:01 | G496S<br>Q498R<br>N501Y | Omicron |
| 4 | QIAPGQTGK | A*68:01 | K417N | Omicron, Beta |
| 5 | SPRRARSV | B*07:02 | P681H | Omicron, Alpha |
| 6 | SPRRARSVA | B*07:02 | P681H | Omicron, Alpha |

**Supplementary Table 2.**
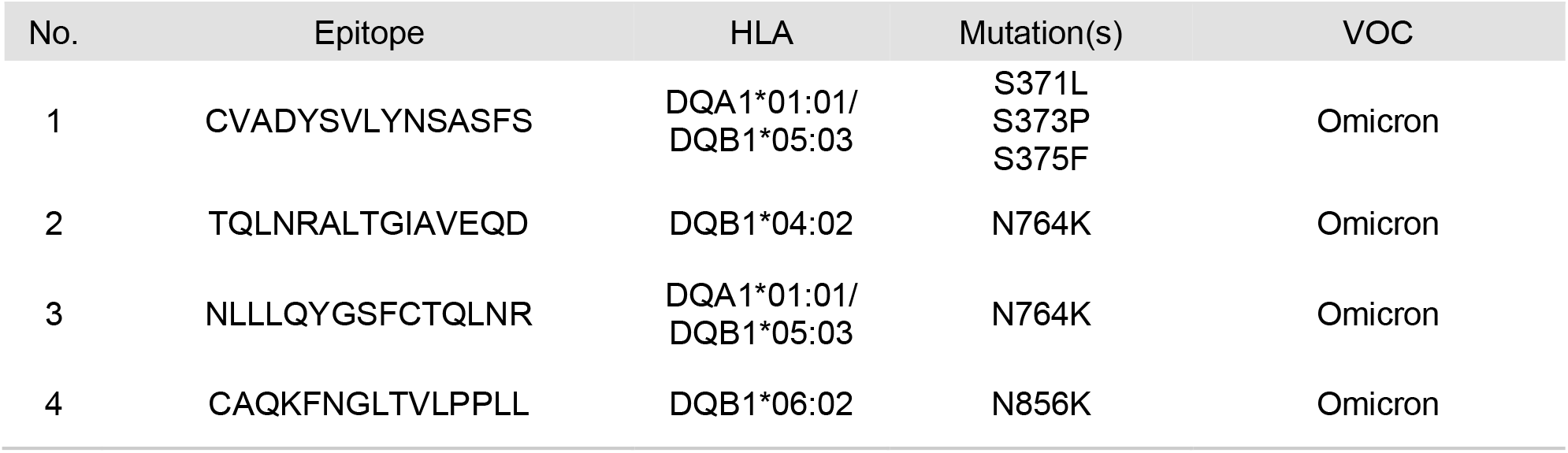
List of CD4^+^ T cell epitopes with Omicron mutations that are predicted to become non-binders. NetMHCpanII-4.0 was employed for predicting peptide-HLA binding using the default parameters (Reynisson et al. 2020).

